# Structural differences between the Woodchuck hepatitis virus core protein in dimer and capsid states indicate entropic and conformational regulation of assembly

**DOI:** 10.1101/512319

**Authors:** Zhongchao Zhao, Joseph Che-Yen Wang, Giovanni Gonzalez-Gutierrez, Balasubramanian Venkatakrishnan, Adam Zlotnick

## Abstract

Hepadnaviruses are hepatotropic enveloped DNA viruses with an icosahedral capsid. Hepatitis B virus (HBV) causes chronic infection in an estimated 240 million people; Woodchuck Hepatitis virus (WHV), an HBV homologue, has been an important model system for drug development. The dimeric capsid protein (Cp) plays multiple functions during the viral life cycle and thus has become an important target for a new generation of antivirals. Purified HBV and WHV Cp spontaneously assemble into 120-dimer capsids. Though they have 65% identity, WHV Cp has error-prone assembly with stronger protein-protein association. We have taken advantage of the differences in assembly to investigate the basis of assembly regulation. We have determined the structures of the WHV capsid to 4.5 Å resolution by cryo-EM and the WHV Cp dimer to 2.9 Å resolution by crystallography and examined the biophysical properties of the dimer. We found, in dimer, the subdomain that makes protein-protein interactions is partially disordered and rotated 21° from its position in capsid. This subdomain is susceptible to proteolysis, consistent with local disorder. These data show there is an entropic cost for assembly that is compensated for by the energetic gain of burying hydrophobic interprotein contacts. We propose a series of stages in assembly that incorporate disorder-to-order transition and structural shifts. WHV assembly shows similar susceptibility to HBV antiviral molecules, suggesting that HBV assembly follows similar transitions and indicating WHV’s importance as a model system for drug development. We suggest that a cascade of structural changes may be a common mechanism for regulating high fidelity capsid assembly in HBV and other viruses.

**Author summary:** Capsid assembly is a key step in the Hepatitis B virus (HBV) life cycle; it requires sophisticated regulation to ensure the production of capsids and viruses with high-fidelity. HBV capsids are constructed from 120 self-assembling capsid protein dimers (Cp). It was hypothesized that changes in Cp structure regulated the assembly reaction. Here we determined structures of dimer and capsid for Woodchuck Hepatitis virus (WHV), an HBV homologue. We observed that the component of the dimer involved in subunit-subunit interactions undergoes a disorder-order transition and changes structure concomitant with assembly. Meanwhile WHV Cp displays similar susceptibility to HBV antiviral. We propose that this structural transition entropically regulates assembly and may be a common theme in other viruses.

## Introduction

Hepatitis B virus (HBV) is an endemic disease that chronically infect around 257 million people. Annually, more than 800,000 deaths result from chronic HBV and its complications, notably liver failure, cirrhosis and hepatocellular carcinoma [1, 2]. Woodchuck hepatitis virus (WHV), a HBV homolog, causes similar symptoms as HBV [3] and has been a critical animal model, contributing to our understanding of HBV and development of antivirals [4, 5].

WHV and HBV have similar architecture; an envelope surrounds an icosahedral capsid, which contains the partially double-strand circular DNA genome [4]. The HBV core protein (Cp), a small 183-residue homodimer, plays multiple roles [6]. Cp dimers specifically assemble around a complex of viral RNA and polymerase. This compartment provides environment for reverse transcription, has signal for nuclear import, has signals for binding viral envelop protein, and releasees mature genomes into the nucleus [7]. Cp dimers are also found in the nucleus associated with viral genomes [8]. The unique characteristics and multiple functions make Cp an important target for developing direct-acting antivirals [6].

HBV capsids have 120 Cp dimers arranged in T=4 icosahedral symmetry [9, 10]. Cp is comprised of a 149-residue assembly domain (hCp149) and a nucleic acid binding domain. The assembly domain has a stable chassis domain linked to sub-domains by glycine/proline hinges (Fig 1a) [11, 12]. *In vitro*, HBV Cp capsid assembly is a reaction with high fidelity [13, 14] that can be initiated and manipulated by pH, ionic strength, temperature using the N-terminal assembly domain of Cp, hCp149 [13]. Hypothetically mis-assembled capsids will lead to failure of HBV lifecycle. *In vitro*studies have provided understanding of the biology of capsid assembly *in vivo*, contributing to development of antivirals that mis-regulate or mis-direct capsid assembly. The Cp of WHV shares 63% sequence identity with HBV Cp [15]. However, the assembly domain of WHV, wCp149, has low assembly fidelity [15, 16]. Kukreja *et al* and Pierson *et al* have shown that wCp149 assembles much faster than hCp149 with stronger protein-protein interactions, frequently leading to irregular and over-sized capsids [15, 16].

**Fig 1.**
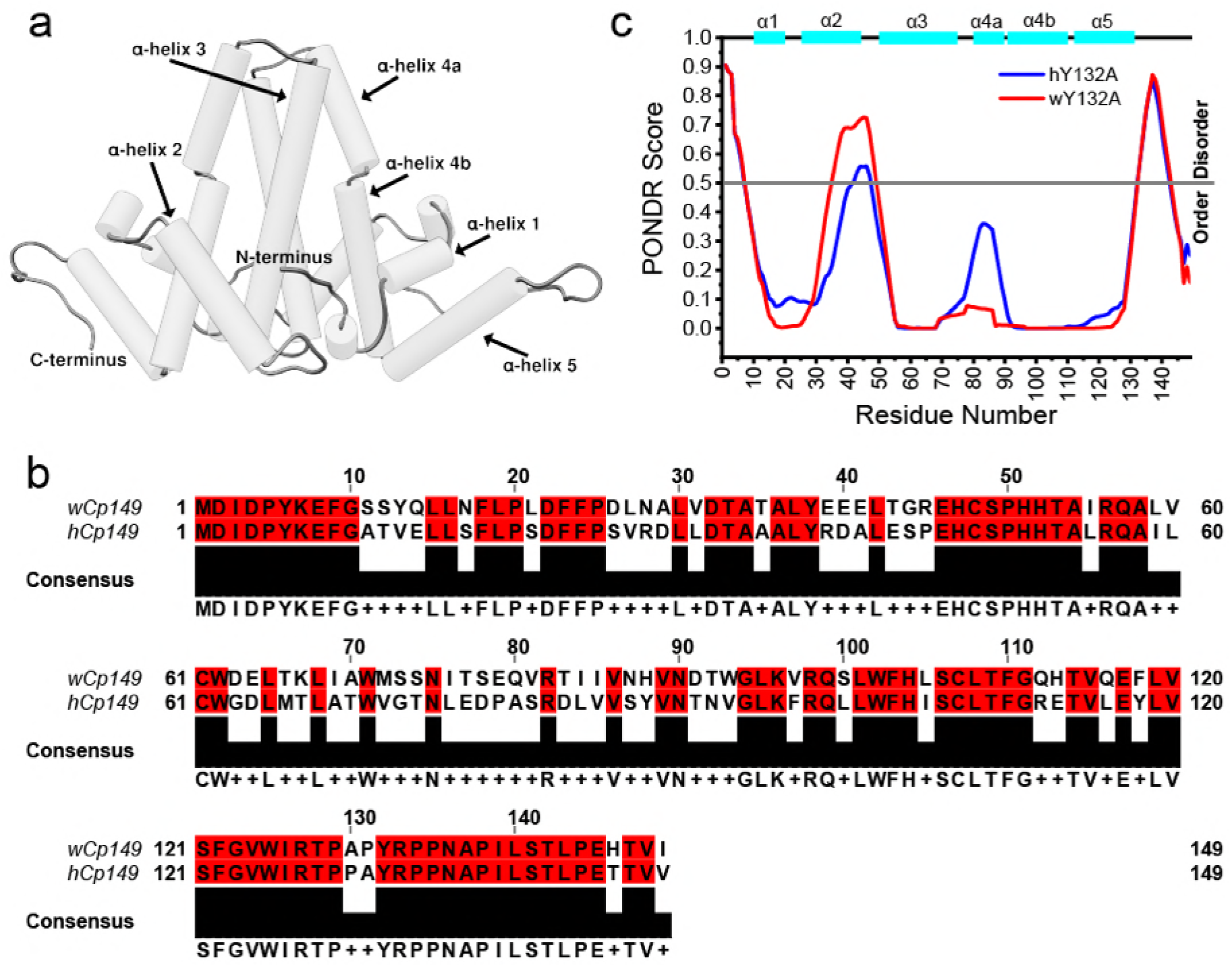
WHV and HBV have a similar assembly domain. (a) A dimer is represented using pipes- and-planks. (b) PONDR VL-XT predicts that hY132A and wY132A have similar disorder pattern specially at the C-termini where missing density is observed (Fig 4c and 4d). (c) Sequence alignment shows that wCp149 and hCp149 have 63% sequence identity.

Here, to identify the structural basis for differences between wCp149 and hCp149, we determined the wCp149 dimer structure using the assembly incompetent mutant [11] wCp149-Y132A (wY132A) and the WHV capsid structure. In the WHV dimer structure, the C-terminal region, residues 110 to 140, that responsible for inter-dimer interaction is shifted by 21° from its location in capsids and is partially disordered. Proteolysis studies confirmed that WHV dimer has a more flexible C-terminus than HBV. Our findings suggest that the entropy of the C-terminal region plays an important role in regulating assembly.

## Results

### wY132A shows unique crystal packing

Kukreja *et al* showed that wCp149 and hCp149 have distinctly different assembly behaviors [15] despite 63% sequence identity (Fig 1b). To understand the basis for those differences, we sought to determine the structure of wCp149 free dimer to compare to hCp149 dimer. Because wild-type wCp149 spontaneously self-assembles, we used an assembly-incompetent mutant, wCp149-Y132A (wY132A), that was designed based on previous work with an assembly incompetent human Cp149-Y132A (hY132) [11, 17]. The Y132A mutation is on an exposed loop of human capsid protein where it is expected to eliminate a substantial fraction of surface that is buried during assembly but have little effect on protein folding. Under various assembly conditions, wY132A showed no assembly based on SEC and light scattering. wY132A did not crystalize under the conditions used to determine the structure of hY132A [11, 12]. After exhaustive screening, we identified and optimized a novel condition for obtaining well-ordered crystals: 13% polyethylene glycol 400, 240 mM KCl and 50 mM MES pH 5.8.

The crystal structure of wY132A was determined to 2.9 Å resolution had two wY132A dimers, AB and CD, in the crystallographic asymmetric unit (ASU) (Fig 2a), in a novel arrangement of dimers. To appreciate the importance of the crystal packing for wY132A, it must be recognized that in crystals of hY132A the protein is always arranged in sheets with triangular repeats (Fig 2c); this arrangement of dimers is based on interdimer contacts that are similar to the interactions found in capsids and involve many of the same protein-protein contacts [11, 12].

**Fig 2.**
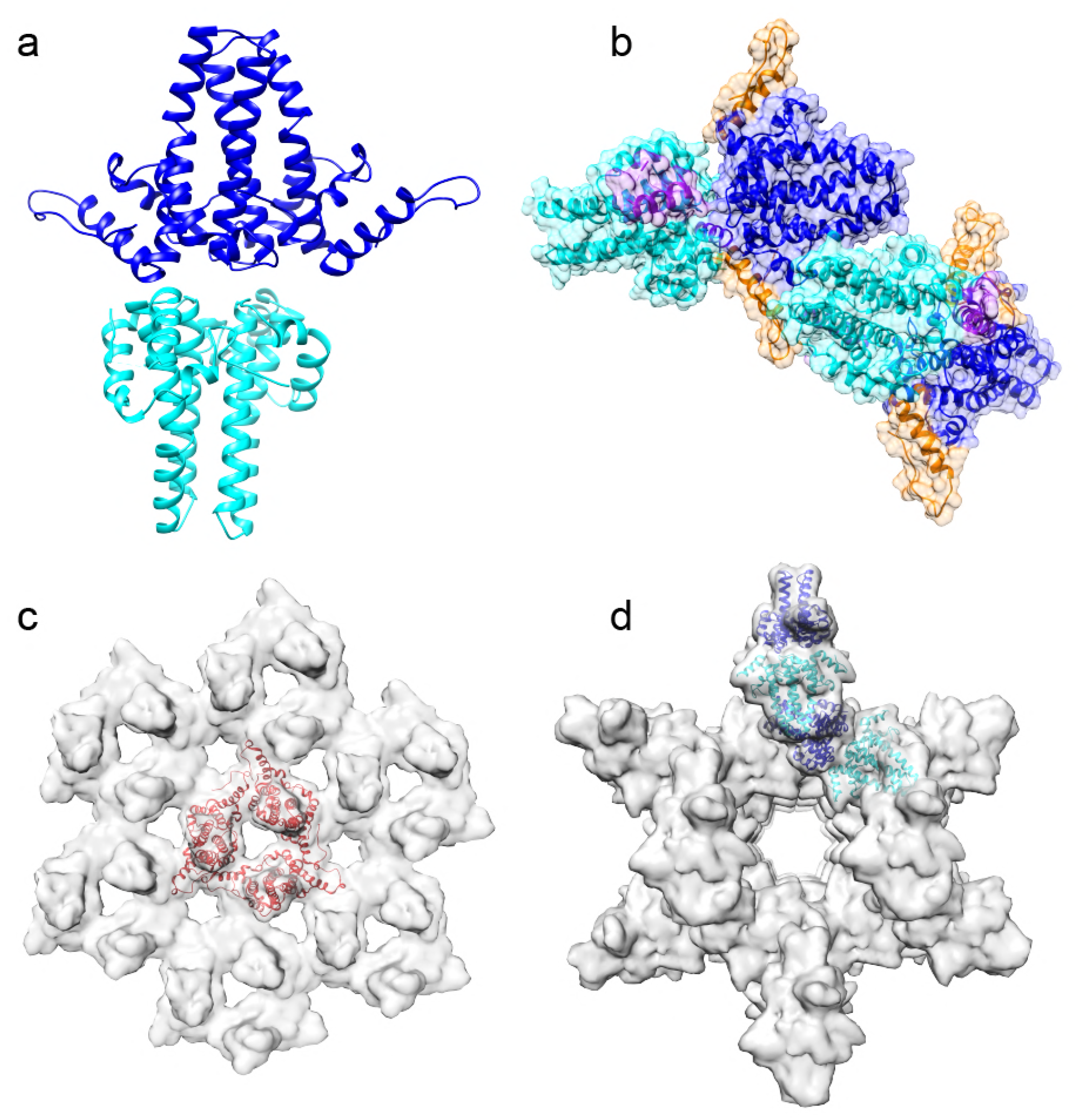
Unlike hY132A crystal packing, wY132A crystal packs differently. (a) The asymmetric unit (ASU) of the wY132A crystal structure have two dimers, AB (blue) and CD (cyan). (b) wY132A repeats its ASU, packing to form a hollow tube. (c) The hY132A (PDB: 4bmg) adopts capsid like interactions to form trimer of dimers (red triangle) when pack form crystal. (d) wY132A interacts differently to form a cylinder in crystal.

In wY132A crystals, interactions between dimers involve contacts not observed in capsids (Fig 2a and 2b). The two dimers, AB and CD, from the wY12A ASU are stacked base to base (Fig 2a), where the base normally forms the lumen of a capsid (Fig S1). ASUs in the crystal are arranged so that the top of the four α-helix bundle of the CD dimer is wedged into the gap formed by α-helix 5 and the four α-helix bundle of the AB dimer from an adjacent ASU (Fig 1a, 2b and 2d). Clusters resulted from such interactions are arranged to form a hollow cylinder (Fig 2d). Crystallization condition did not cause wY132A to form cylinders in solution based on TEM, suggesting that such interaction and structure are specific for crystal packing.

### wY132A dimers adopt a single conformation but with some importance differences

Though the two dimers in the crystallographic asymmetric unit have very different environments, thy have nearly the same structure, excepting some missing density in the CD dimer. In overlays, all monomers in the wY132A asymmetric unit had one conformation: A, B, C, and D aligned with little deviation, despite the missing density in C and D (Fig 3c, 3d, 4c and 4d). The RMSD of the alignments were 0.37 Å (AB), 0.67 Å (AC) and 0.53 Å (AD). In contrast, a T=4 HBV capsid [9] has two asymmetric dimers resulting in four unique monomer structures and the hY 132A 3KXS crystal structure [11] has three asymmetric dimers resulting in two classes of monomer structures. The wY132A dimers nearly have a two-fold symmetry off by only 0.65°; AB and CD dimers of wY132A aligned with little difference in Ca position over both subunits and both dimers (Fig 3).

**Fig 3.**
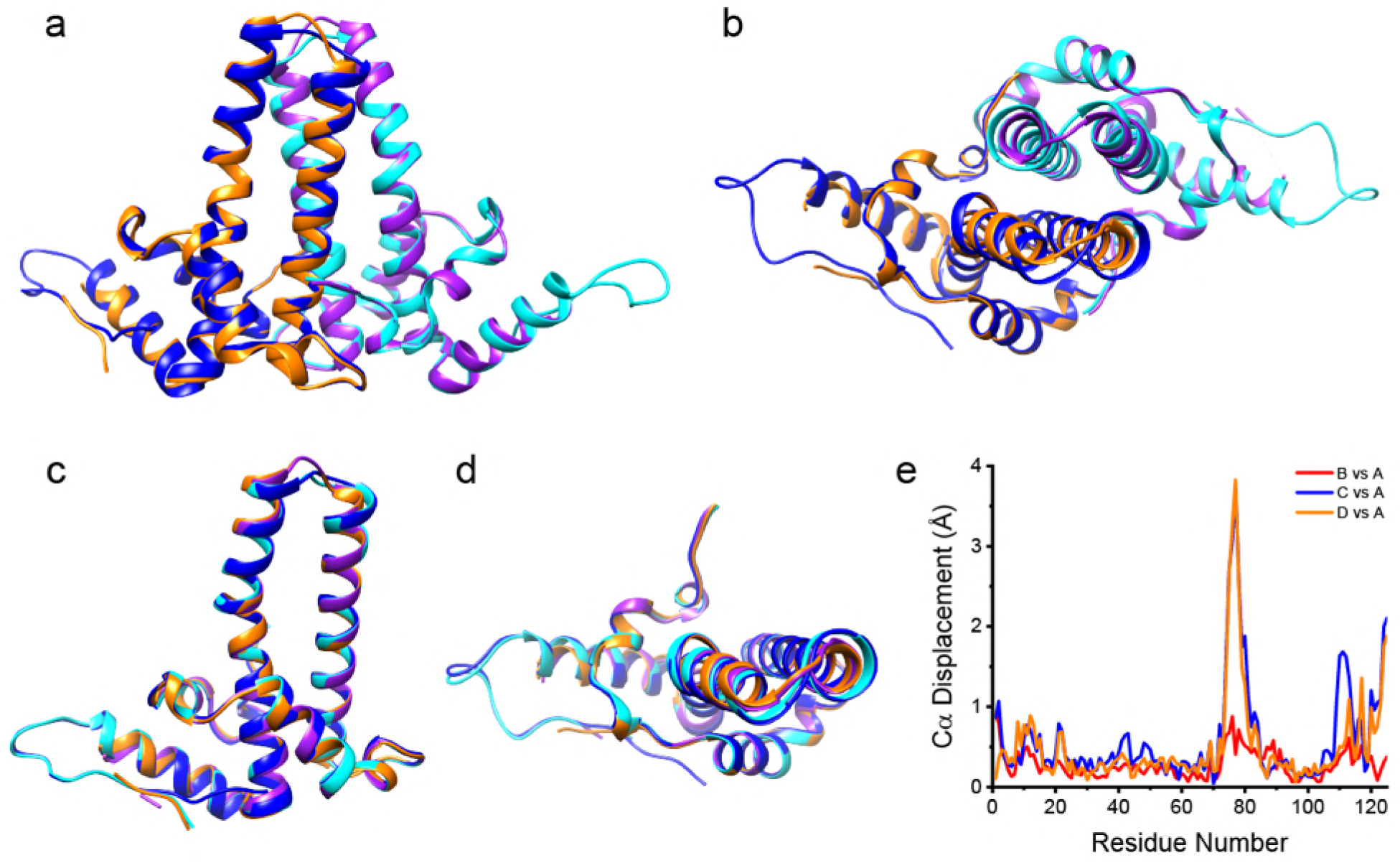
All subunits from the wY132A crystal structure asymmetric unit adopt one conformation. (a, side view and b, top view) Structure alignment of AB dimer and CD dimer shows that both dimers have similar tertiary structures. (c, side view and d, top view) Overlapping all subunits (A, B, C and D) shows that all subunits from the asymmetric unit exhibit a single conformation. (e) To quantitively compare four subunits, the Cα displacements of chain B, C and D comparing with chain A is calculated and plotted against residue number. As shown in (d), most of the residues has displacement lower than 1 Å, indicating that all four subunits overall have a similar tertiary structure except regions (the spike tips and C-termini) where electron density starts to degrade.

Despite the overall conformational similarity, the AB and CD dimers display an important difference in their density maps (Fig 4). At 1.5 σ contour level, the electron density map of the AB dimer is continuous, with no breaks (Fig 4a and 4b). However, at the same contour level, the map for the CD dimer has missing density corresponding to residues 126 to 136, a loop structure linking α-helix 5 to the C-terminal extended structure of the assembly domain (Fig 4c and 4d). This disorder has never been observed in hCp149 capsid or hY132A dimer structures. The ordered density in the loop of the AB dimer corresponds to packing interactions with neighboring CD dimers. The equivalent loop in CD dimers is surrounded by solvent. The missing density of the loop in CD dimers leads us to suggest that the loop is similarly disordered when the dimer is in solution. Though this region is not involved in crystallographic contacts for CD dimers, it plays a critical role in the dimer-dimer interactions involved in building capsids [9, 10]. The disorder of the sequence from residue 126 to 136 is consistent with the proteolytic sensitivity of Arg127 in human HBV Cp149 [18].

**Fig 4.**
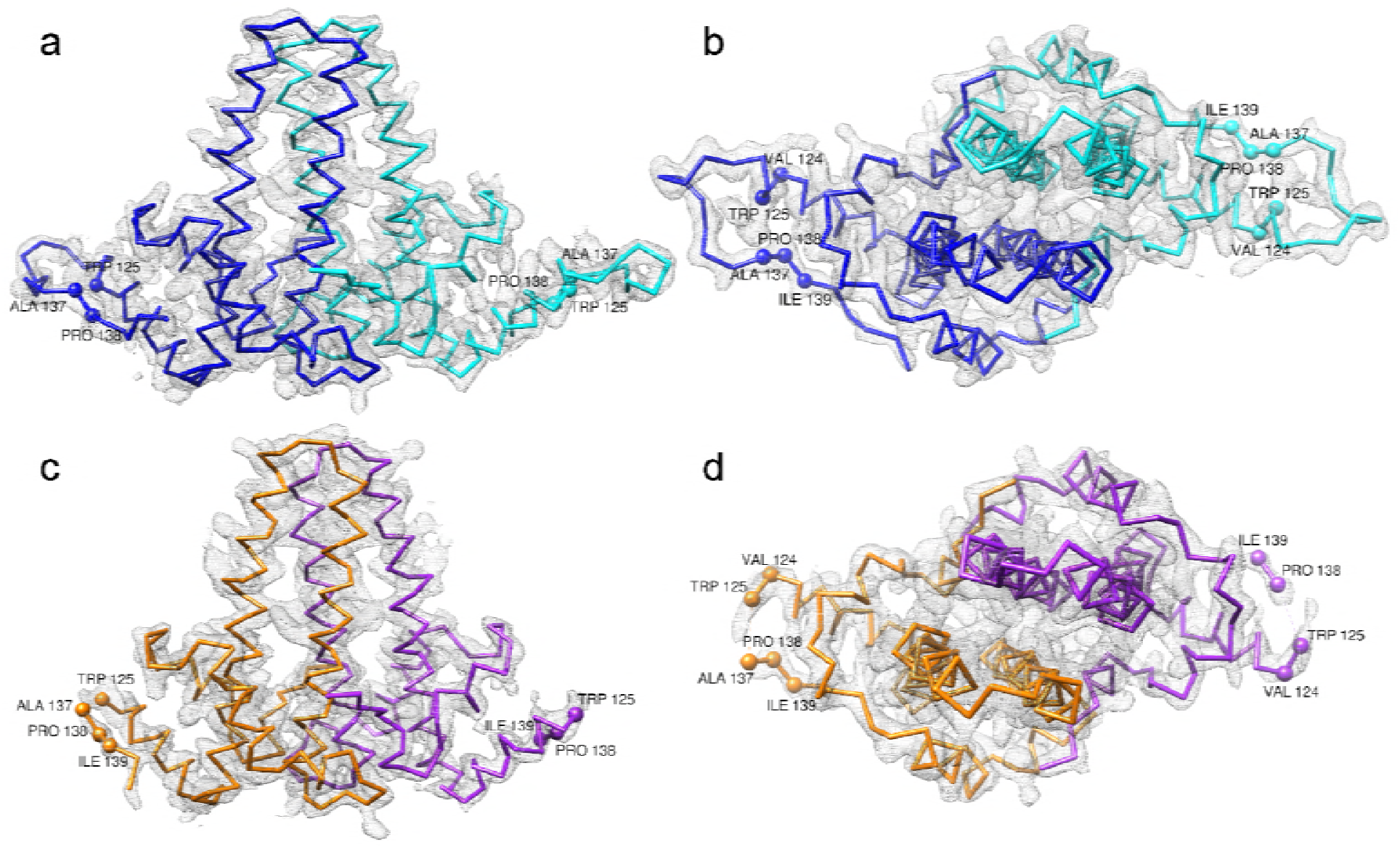
wY132A dimers have different features. 2Fo-Fc electron density map of wY132A crystal structure at 1.5 σ contour level with Cα chains and its subunits comparison among an asymmetric unit, with A subunit in blue, B subunit in cyan, C subunit in orange and D subunit in purple. (a) Side view and (b) top view of AB dimer show that electron density covers the entire backbone. In comparison to AB dimer, (c) side view and (d) top view of CD dimer show missing electron density at the loop turn of C-termini, roughly from residue 126 to 136, where dimers make contacts during assembly. Residues are labelled to show the missing part. The molecular model has PDB accession code 6ECS.

### wY132A and hY132A have substantive structural and biochemical differences

Human HBV dimers have a central core with mobile peripheral sub-domains [9, 11]. This also seems to be the case with wY132A dimers (Fig 2a, 3a, 3b and 4) but with differences. The AB dimer of wY132A was superimposed with the EF dimer (PDB: 3KXS), a 2.25 Å resolution structure of hY132A (Fig 5). There was a large difference in Cα positions at the spike tips connecting α-helix 3 and α-helix 4a, a notably flexible region [6]. The top view of the overlay also showed that α-helix 5 of wY 132A is rotated about 8.5° away from hY132A, using residue Gly111 as a hinge residue (Fig 5b). With this position for α-helix 5, wY132A could not pack into trimer of dimers as seen in the crystallographic packing of hY132A (Fig 2c) [11]. Comparisons of the two aligned structures showed that the Cα positions for the spike tips (residues 60 to 100) and the α-helix 5 inter-dimer interaction region were displaced up to 8 Å whereas core regions are displaced by less than 1 Å (Fig 5c).

**Fig 5.**
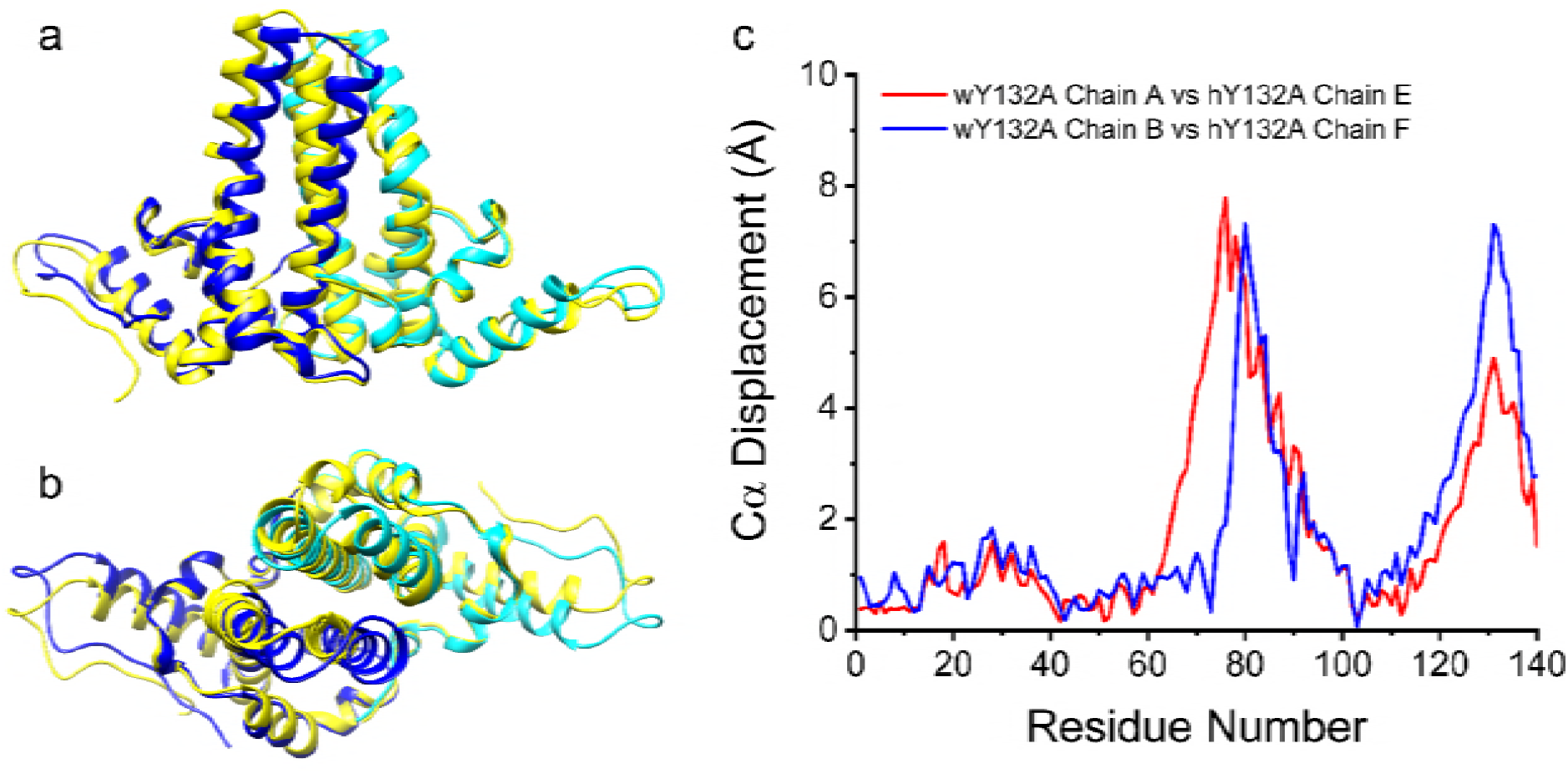
wY132A AB dimer has different tertiary structures than hY132A. (a and b) wY 132A AB dimer is overlaid with hY 132A EF dimer (labelled in yellow). The top half of the four α-helix bundles and C-termini show distinct structure shift, which is quantified by plots Cα displacements against residue numbers (c).

To test if the sequence of the disordered loop in wY132A and the equivalent region of hY132A had a predilection for disorder, we used PONDR (http://www.pondr.com/) to analyze both proteins. Given the 63% sequence identity (Fig 1b), it is not surprising that the two proteins led to similar predictions (Fig 1c). Both sequences had a notable peak in predicted disorder score at the residues 126-136 loop as well as a region from residues 35-50 that interacts with helix 5, which had been suggested as a fulcrum that modulated α-helix 5 mobility [11].

To investigate whether disorder in the loop from residues 126 to 136 of wY132A corresponded to increased protein flexibility compared to hY132A, we tested their susceptibility to proteolysis. In hCp149, it was discovered that Arg127, though part of α-helix 5, was the first site cleaved by trypsin [18]. In the current study, we observed that wY132A was digested by trypsin much faster than hY132A (Fig 6a). As expected from previous work [18], cleavage of both proteins was at Arg127, confirmed by mass spectrometry. In the earlier study, we determined that unfolding, or “opening”, of the Arg127 site was the rate limiting factor for proteolysis. The faster digestion of wY132A suggests residues 126-136 are more disordered. To test if the faster cleavage rate of wY 132A indicates that the protein is partially unfolded in solution and had a correspondingly larger hydrodynamic radius, we used size exclusion chromatography (SEC). Paradoxically, wY132A eluted earlier than hY132A, suggesting that wY132A has a more compact Stokes’ radius (Fig 6b).

**Fig 6.**
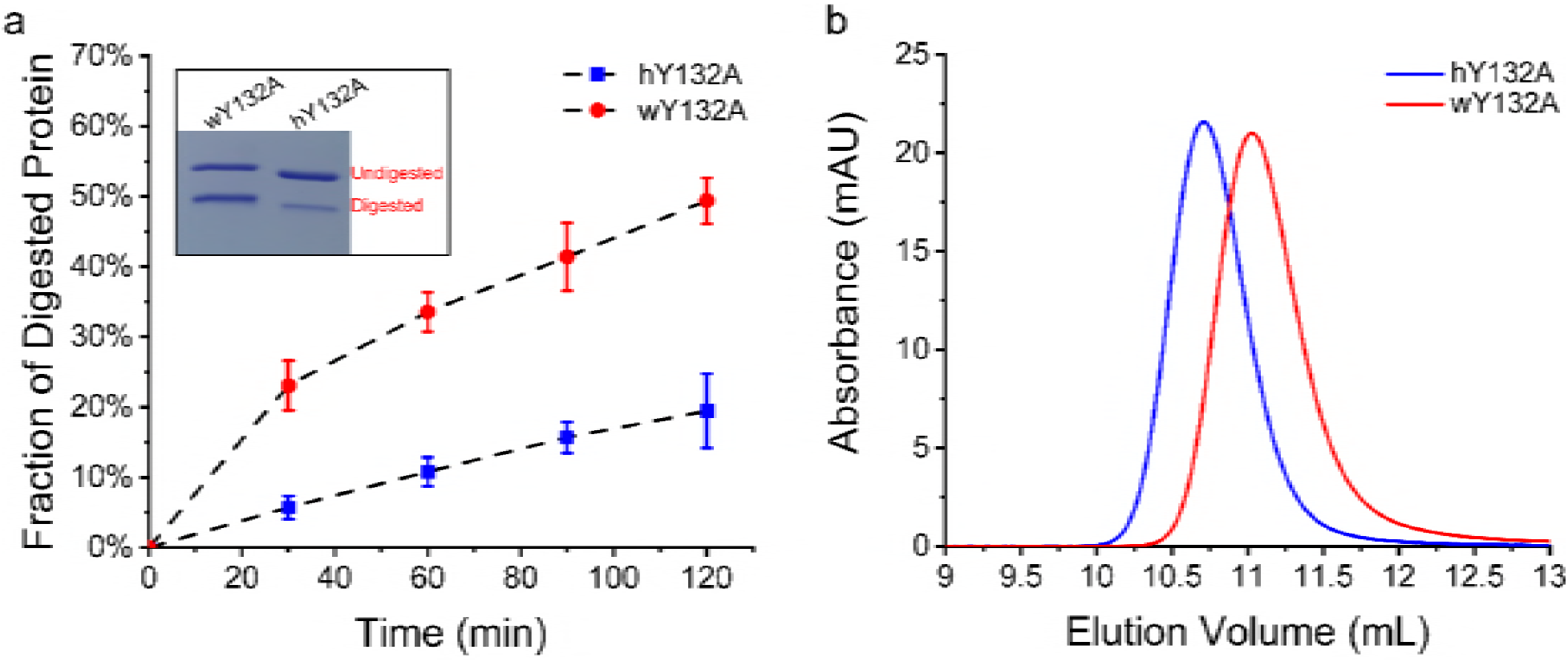
hY132A and wY132A have different rates of proteolysis in a region where the sequence is identical. (a) Proteolysis of hY132A and wY132A by trypsin is quantified by SDS-PAGE and shows that wY132A (red dot) is digested much faster than hY132A (blue square). The inset is an example of SDS-PAGE of trypsin digestion. (b) An SEC experiment of hY132A and wY132A with high salt (1 M NaCl) shows that wY132A elutes later than hY132A, indicating that wY132A has a smaller Stokes radius and a more compact form.

### wCp149 from dimer to capsid: assembly is more thermodynamically favored than hCp149

Assembly of hCp149 and wCp149 are temperature-dependent and entropy-driven reactions, consistent with the observed burial of hydrophobic surface at the dimer-dimer contacts [6]. However, wCp149 dimers have stronger association energies and its assembly is favored in a broad range of temperature [15]. To quantify assembly of wCp149, we used very low ionic strength (50 mM NaCl and 10 mM HEPES pH 7.5) at different temperatures, conditions where hCp149 showed no assembly (Fig S2a). We noticed that under these assembly conditions, wCp149 has a linear van’t Hoff plot, which suggested that wCp149 assembly has a close to zero heat capacity (Fig S2b). This is different from the positive heat capacity previously obtained using 100 mM NaCl [15].

### wY132A needs conformational changes to form capsid

Superposition of wY132A AB dimer with AB and CD dimers of HBV capsid (PDB: 1QGT) revealed a critical structure difference in the position of α-helix 5, which rotated 21° from the dimer in capsid form (Fig S3 and S4). Even though AB and CD dimers of HBV capsid are asymmetric dimers, they display a similar pattern of conformational change compared to wY132A (Fig S3 and S4). In addition to the highly variable spike tips (residue 80 to 90), Cα positions for the region containing residues 120 to140 move up to 12 Å (Fig S3 and S4).

To test whether WHV core proteins have a different conformation in the capsid form or if the dimers change their conformation when they assemble, we determined the cryo-EM structure of WHV capsid to 4.5 Å resolution (Fig 7a and 7b). Comparison between WHV (Fig 7a) and HBV (Fig 7c) capsid density map suggests that they are virtually identical (Fig 7d). This observation led us to infer that wCp149 dimers undergo structural changes during assembly. Furthermore, the structural similarity of WHV and HBV indicates that WHV is suitable model system for testing and development of capsid-directed antivirals.

**Fig 7.**
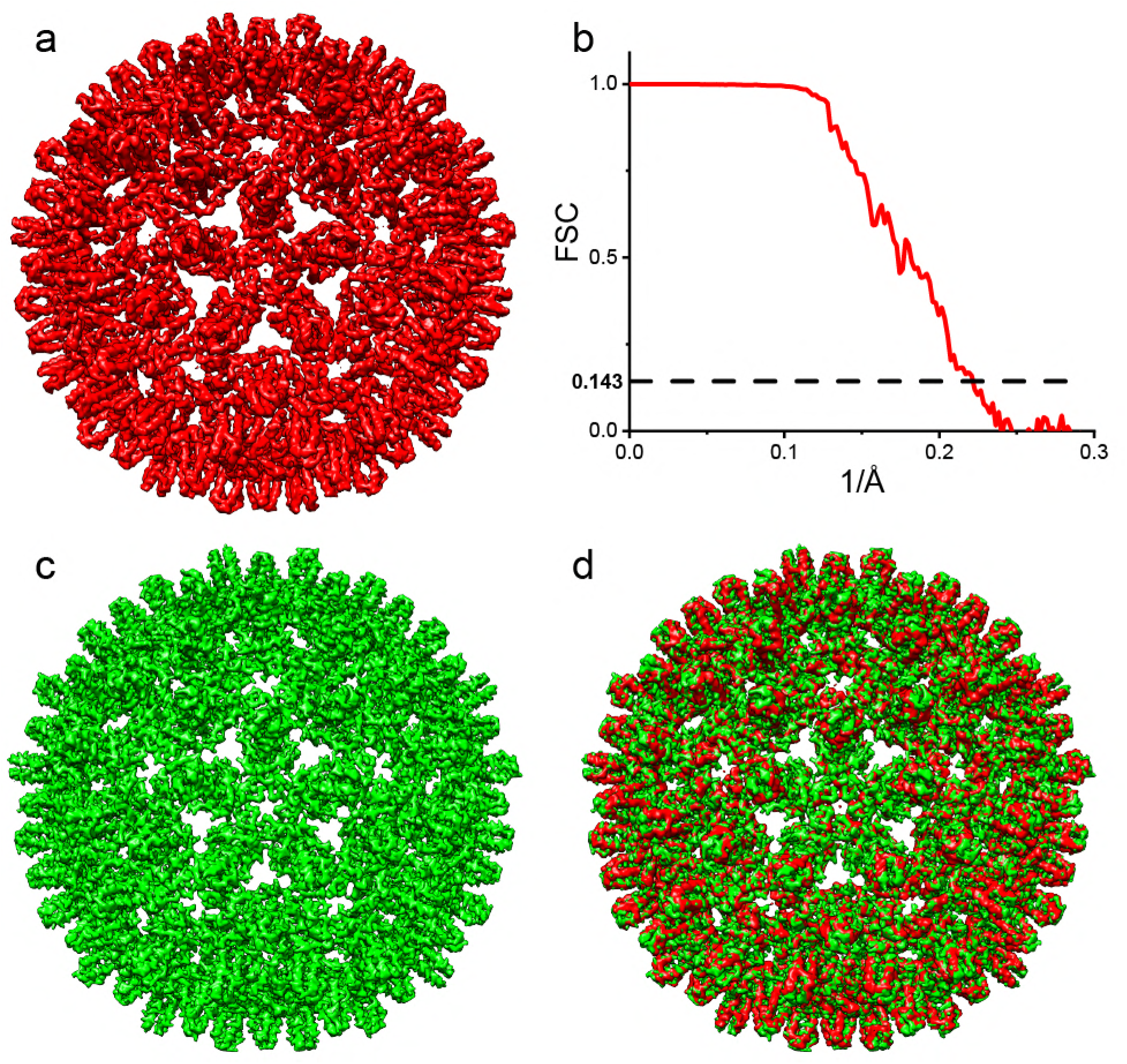
WHV and HBV have similar Structures. Cryo-EM density map of WHV capsid (a) and simulated HBV capsid density map (c) show icosahedral symmetry. Overlapping WHV and HBV capsid density (d) shows that they have similar capsid structures. (b) Fourier shell correlation for the WHV capsid reconstruction. The dashed line indicates a correlation of 0.143. The density map has EMDB accession code 9031; the resulting molecular model has PDB accession code 6EDJ.

To prove this hypothesis, we further compared the wY132A crystallographic dimer model to the cryo-EM WHV capsid density: a quasi-sixfold hexamer extracted from the WHV capsid to avoid biased interpretation. Six wY132A dimers were fit as rigid bodies into the extracted capsid density, a process that was dominated by the successful match of the four-helix bundle at the dimer interface (Fig 8). The majority of the wY132A molecular model matches the density map except the C-terminal from helix 5 to the end of the visible protein density, residues 112 to 141 (Fig 8). Dimers with residue alanine 132 reverted back tyrosine were refined to density using molecular dynamics flexible fitting (MDFF) [19] and PHENIX real space refinement with icosahedral symmetry constraints [20]. The resulting models fit into the hexamer density (Fig 8). These models are nearly identical to the corresponding dimers from human HBV capsid [9]. Nevertheless, comparing the wY132A dimers and dimers from the refined WHV capsid structure shows major changes at the regions following Gly111 and modest changes at the spike tip and α-helix 2 (Fig S3 and S4).

**Fig 8.**
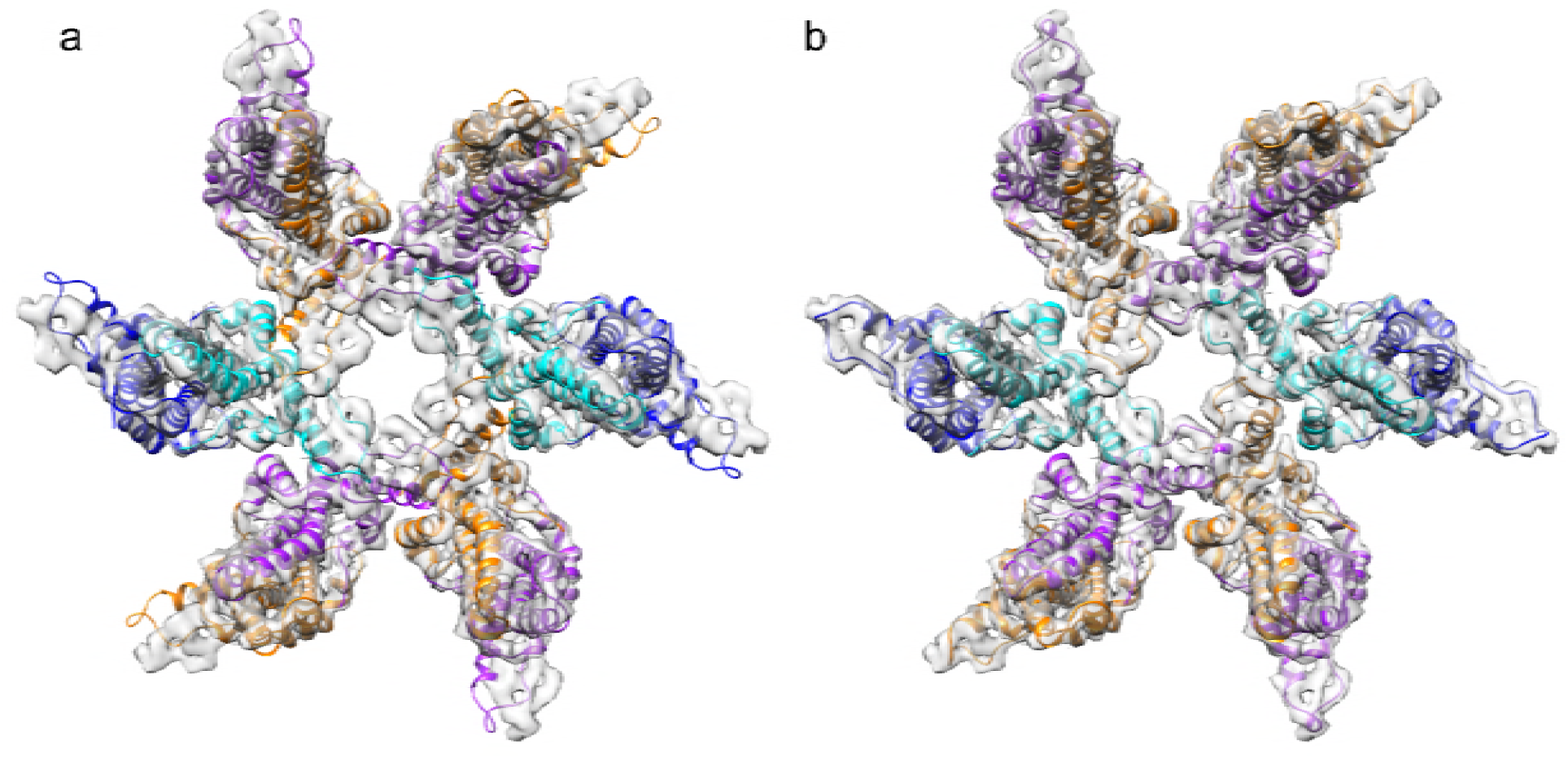
wY132A AB dimers did not fit into WHV capsid density. (a) wY132A AB dimers were rigid body fitted into WHV hexamer density. The crystal structures and the capsid density map match each other well, but with the C-termini as exceptions. All the C-termini toward outside are out of the density, and the C-termini located inside intrude into the adjacent subunits, causing clashes. (b) Same capsid hexamer density map was occupied by determined WHV capsid structure, showing the regions for structural moving.

### wCp149 assembly can be distorted by assembly-directed antivirals

WHV and human HBV have similar capsid structures (Fig 7), supporting the feasibility of WHV as a model system for testing and developing assembly-directed antivirals. To test this hypothesis, wCp149 assembly was observed with or without two antiviral molecules, HAP12 and HAP13 [21], and their assembly products were observed using TEM. Under conditions where wCp149 showed modest assembly, HAP12 significantly increased wCp149 assembly and HAP13 showed the similar assembly effect but to a lesser extent (Fig 9a). wCp149 mainly assembles into T=4 capsids at physiological ionic strength (150 mM NaCl pH 7.5) (Fig 9b). With the addition of HAP12, wCp149 assembles into products with larger, aberrant complexes, often with large openings (Fig 9c and 9e). Consistent with its effect on assembly kinetics, HAP13 led to aberrant assembly products, but defects were not as gross as with HAP12 (Fig 9d and 9f). These observations for wCp149 parallel our previous results with HBV Cp149 [21], supporting the utility of using WHV as a tool for testing and developing assembly-directed antivirals.

**Fig 9.**
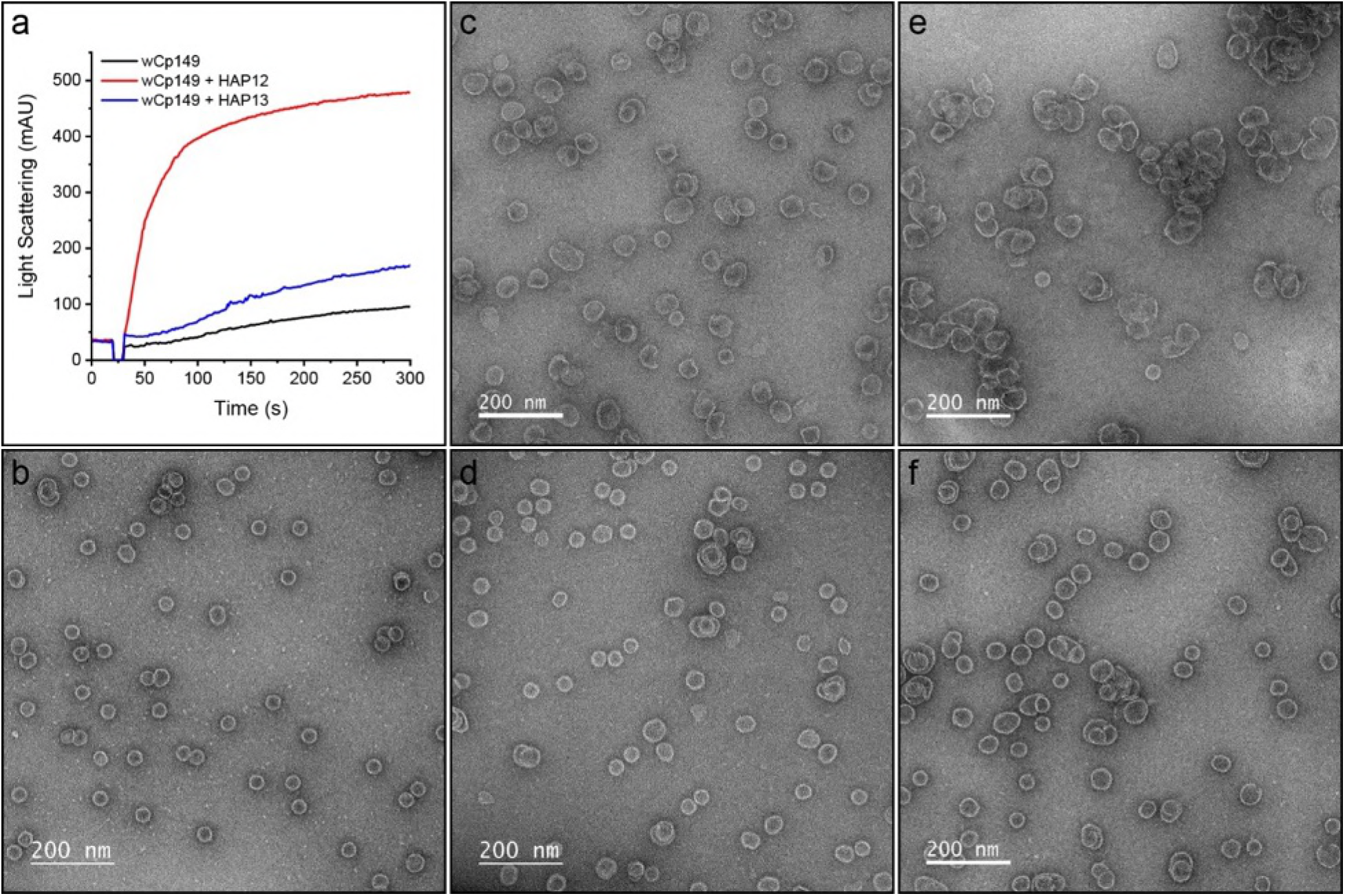
wCp149 assembly is distorted by antiviral molecules, HAP12 and HAP13. (a) Under conditions where wCp149 has slight assembly, 4 μM HAP12 and HAP13 increased wCp149 assembly. (b) In the absence of HAPs, most of the wCp149 assembly products are circular T=4 capsids. With the addition of 4 μM HAP12 (c) or HAP13 (d), wCp149 assembles into a mixture of oversized products with few normal-sized capsids. Increasing the antiviral concentration to 10 μM (e, HAP12; f, HAP13) showed more dramatic effects on the assembly products. These negative stained micrographs were recorded at 50,000x magnification.

## Discussion

To investigate the basis of regulated HBV capsid assembly, we have determined the structures of wY132A and a WHV capsid. The wY132A structure is nearly symmetric and the dimers are not involved in capsid-like interactions (seen in hY132A structures [11, 12]). Observed wY132A symmetry is consistent with NMR analysis of hCp149, albeit at pH 9.5 [22]. Without the constraint of quaternary structure in the crystallographic packing, the subdomain comprised by α-helix 5 and following residues pivot around Gly111, shifting 8.5° away from its position in hY132A, which is constrained by a trigonal crystallographic lattice and 21° from hCp149 in a T=4 capsid (Fig 5, 10 and S3 and S4 Fig). Though the two wY132A dimers in the ASU are nearly identical, the CD dimer has missing density for residues 126-136, whereas those residues are ordered in AB dimer (Fig 4). The missing density indicates that in the WHV dimer, this disordered loop fits a large ensemble of conformations. The disorder region matches PONDR predictions (Fig 1c) and the previously observed susceptibility of Arg127 to proteolysis [18]. In this study, we observed a much faster digestion rate for wY132A (Fig 6a), suggesting that in WHV dimers, this loop is more dynamic. Structurally disordered regions have been proposed to possess functional roles, becoming ordered upon interaction with a target molecule or protein [23, 24]. In WHV Cp, the disordered loop becomes ordered by interaction with another dimer during capsid assembly. Although there is an entropic cost for ordering the loop, it must be offset by the energetic advantage of assembly. We suggest that wY132A structure, with its symmetry, shifted α-helix 5, and the disordered loop, is an instructive representative of free dimer.

In previous work, we observed that the association energy of assembly was less than predicted based on structure and suggested that the difference was the energetic cost of a conformational change relating to assembly [11, 13]. This hypothesis is qualitatively consistent with the shift of α-helix 5 (residues 111-128) and ordering of the associated loop (residues 136-136). Similarly, molecular dynamics simulations show that even in a stable HBV capsid, dimers change conformations constantly [25]. This leads us to propose two models for regulation of capsid assembly.

The first model of assembly is based on sequential conformational change. We have observed three distinct conformations for dimer: (i) the partially disordered form shown here for wY132A, (ii) planar sheets of triangles found in hY132A crystals, and (iii) capsids where dimer-dimer interactions conform to a capsid’s radius of curvature. We suggest there three conformations corresponds to free dimer, early assembly intermediates before curvature is locked in, and capsid-like assembly intermediates and products. Assembly factors (i.e. temperature, ionic strength, and pH), trigger a conformational change by rotating α-helix 5 by 8.5° so that free dimers can associate into triangles to form nuclei and small oligomers. As the assembly products grows, the planar sheet curves forcing further rotation of α-helix 5 to 21° which then leads to T=4 icosahedral capsids (Fig 10).

**Fig 10.**
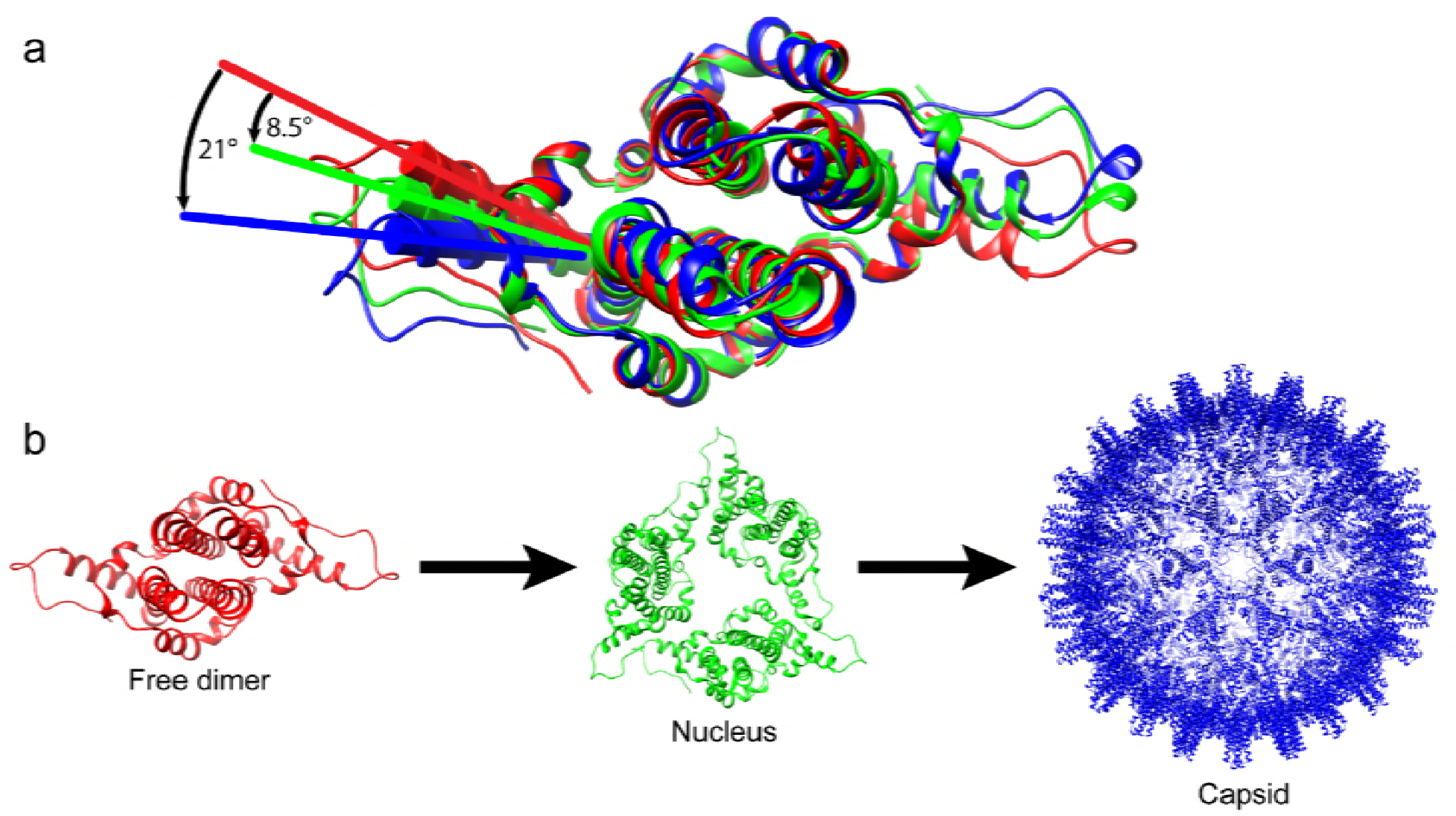
α-helix 5 modulates the capsid assembly process by changing its conformations. (a) Alignment of wY132A (red), hY132A (green, PDB:3kxs) and dimer in WHV capsid (blue) shows that three dimers have different α-helix 5 and C-termini. wY132A needs to shift 8.5° to match the conformation of hY132A and 21° to wCp149. (b) A scheme of assembly process under different dimer conformations (Structure labeled using same colors corresponding to Fig 10a). Assembly starts with free dimers (wY132A labelled in red). Initiation of assembly induces around 8.5° shift of α-helix 5 to generate nuclei (hY 132A, trimer of dimers) [11, 13]. As assembly proceeds, the α-helix of dimers keep shifting to the extent of 21°, under which dimers have the correct conformations to interact with each to form stable capsids (WHV capsid in blue).

The second model posits that assembly is regulated entropically [23]. In the entropic model, dimers have an ensemble of conformations that changes in response to solution conditions. For free dimers, this presumably includes equilibration between ordered and disordered states for the loop from residues 126-136. As stated by Hilser *et al.*, the effect of allosteric coupling between two disordered regions will accelerate the conformational changes and dimer-dimer interactions because binding of one site of disordered region will cooperatively turn the other disordered region into ordered form and increase the binding affinity to new incoming subunits [23]. Therefore, the energetic cost of ordering the first few subunits provides a basis for the nucleation phase. The allosteric effect of subsequent assembly suggests an element of cooperativity (Fig 10), an idea originally described as autostery by Caspar [26]. Importantly, these models are not mutually exclusive.

Both models provide explanations for normal and abnormal assembly reactions. Because of the presence of different conformations under different assembly stages, there must be an orchestrated assembly process to adjust the protein conformations to produce high-fidelity capsids. Conversely, random shifts of α-helix 5 position will almost certainly lead to incorrect dimer-dimer geometry, eventually forming an abnormal capsid. This effect has been demonstrated with assembly induced by small molecules that bind to a pocket formed by α-helix 5, residues 136-140, and α-helix 2 [10].

The structural similarities of WHV and HBV capsids suggest that human HBV undergoes the same transitions observed for WHV. These similarities extend to the WHV and HBV responses to two core protein-directed antiviral compounds, HAP12 and HAP13 (Fig 9). With human HBV, HAP12 had a much greater effect than HAP13 on assembly and antiviral activity [21, 27]. Similarly, HAP12 had a stronger effect on wCp149 than HAP13, though both led Cp149 and wCp149 assembly to form complexes that were oversized, elongated and even broken (e.g. Fig 9). Of note, the effect of HAP13 on WHV is attenuated compared to what it does to HBV (see Fig 5 in Ruan et al [27]). We suggest that this difference arises because wCp149 assembly association energy is stronger than that of Cp149 (Fig S2); similarly, it was observed that core protein mutations that stabilize HBV capsids confer greater resistance against the weaker HAPs [27]. This set of observations is also consistent with the hypothesis that, in the presence of assembly-directed antivirals, there is a completion between normal and aberrant assembly [28].

In conclusion, we observed that dimers have different conformations at different assembly stages. Loss of disorder in the 126-136 loop, as well as conformational shift of α-helix 5, suggests there is an entropic cost for assembly in addition to the entropic gain from burial of hydrophobic surface. We propose that regulating conformation, or ensemble of conformations, can trigger assembly and control the cascade of assembly reactions that produce uniform 120-dimer T=4 HBV capsids.

## Materials and Methods

### Cloning of wCp149-Y132A (wY132A) and proteins purification

The Woodchuck Hepatitis virus (WHV) capsid protein assembly domain (wCp149) was expressed and purified as described previously [15]. The expression plasmid for wCp149, pET11c-wCp149, was mutated to wCp149-Y132A (wY132A) using QuikChange mutagenesis (Stratagene). wY132A and its human homolog hCp149-Y132A (hY132A) were expressed and purified using the same previously described protocol [11, 17].

### Protein crystallization

wY132A was dialyzed into 50 mM Tris pH 9 and concentrated to 17.5 mg/mL using an Amicon Ultra 30 kDa centrifugal filter. A crystallization screening of a variety of conditions gave a promising hit with 12% polyethylene glycol 400, 400 mM KCl, and 50 mM HEPES pH 7.5 at room temperature. Crystallization conditions were refined to 13-16% polyethylene glycol 400, 220-320 mM KCl, and 50 mM MES pH 5.8 in a sitting drop format. The crystal used for solving the wY132A structure grew in 13% polyethylene glycol 400, 240 mM KCl and 50 mM MES pH 5.8. For data collection, crystals were cryoprotected by briefly soaking in 25% glycerol, 15% polyethylene glycol 400, 260 mM KCl, and 50 mM MES pH 5.5.

### wY132A structure determination and refinement

Diffraction data were collected at the Advanced Light Source (ALS), beamline 4.2.2. Data were processed using the XDS package (XDS, XSCALE and XDSCONV) [29]. The high resolution cutoff was set to 2.9 Å to ensure that values of I/σ and CC_1/2_ in the highest resolution bin (3.0-2.9 Å) were higher than 1 and 0.5, respectively [30]. A molecular replacement density map was calculated using a dimer from the hY 132A (PDB: 3KXS) [11] as the phasing model. Iterative refinement and manual model building were accomplished with programs in the Phenix suite [31] and Coot [32]. TLS (Translation/Libration/Screw) refinement notably improved the R factors. Throughout the refinement process, 7% of the reflections were used for R_free_ to avoid structure overfitting. For the final model, R_work_ was 19.8% and R_free_ was 22.8% (S1 Table). The final structure coordinates of wY132A are deposited in the protein data bank as 6ECS.

### Proteolysis reactions

hY132A and wY132A were first dialyzed into 50 mM HEPES pH 7.5 at 4°C. Sequencing-grade modified trypsin (Promaga) was stored as an inactive stock in 50 mM acetic acid at a concentration of ~4 μM. Each proteolysis reaction consisted of 10 μM hY132A or wY132A, 0.2 μM trypsin and 50 mM HEPES pH 7.5 in a total volume of 200 μL. At time points 0, 30, 60, 90 and 120 mins, 10 μL of each reaction was mixed with 10 μL of denaturing loading buffer and boiled for 7 mins.

### Solution characterization of dimers and dimer assembly

Dimer hydrodynamic radii were compared by size-exclusion chromatography (SEC) using a Superdex 75 10/300 GL column (GE Healthcare) equilibrated with 1 M NaCl 50 mM HEPES pH 7.5. hY132A and wY132A were dialyzed into 50 mM HEPES pH 7.5 at 4°C and adjusted to 5 μM. Proteins were mixed 1:1 (V:V) with 2 M NaCl and 50 mM HEPES pH 7.5 and incubated overnight at room temperature.

To characterize the thermodynamics of wCp149 assembly, assembly reactions were equilibrated, and the amounts of capsid and dimer were determined by SEC. As previously published [15], 10 μM wCp149 was mixed with 50 mM NaCl 10 mM HEPES pH 7.5 and incubated for 48 hours at temperature 9°C, 20°C and 30°C using a Superose 6 10/300 GL column.

wCp149 assembly kinetics were characterized using 90° light scattering as described previously [13]. For control reactions, 4 μM wCp149 was mixed with an equal volume of 300 mM NaCl 10 mM HEPES pH 7.5, giving 2 μM wCp149 150 mM NaCl 10 mM HEPES pH 7.5. For reactions containing antivirals, 8 μM or 20 μM HAP12 or HAP13 in 1% DMSO were mixed with the salt solution (300 mM NaCl 10 mM HEPES pH 7.5); then, equal volumes of protein and salt were mixed, giving reactions with 4 μM or 10 μM HAP12 or HAP13. To visualize assembly products by TEM, 4 μL of each assembly reaction was applied to a glow-discharged carbon film 300 mesh Cu grid and stained with 2% uranyl acetate. Micrographs were collected using a JEOL JEM 1400plus transmission electron microscope equipped with a 4k x 4k CCD camera.

### Cryo-EM structure determination of WHV capsids

wCp149 was dialyzed into 50 mM HEPES pH 7.5 at 4°C and 25 μM wCp149 was assembled by mixing 1:1 (V:V) with 300 mM NaCl, 50 mM HEPES pH 7.5 and incubated overnight. The wCp149 assembly mixture was loaded onto a 10-40% (w/v) continuous sucrose gradient containing 300 mM NaCl, 50 mM HEPES pH 7.5 and centrifuged at 285,000 g for 5 hours in a SW40 swinging bucket rotor. wCp149 capsids, visible as well-defined bands, were extracted from the gradient tube, dialyzed into 300 mM NaCl, 50 mM HEPES pH 7.5 and concentrated to 15 mg/mL.

Preparation of wCp149 capsids for Cryo-EM and image analysis were followed previously described methods [10]. Briefly, 4 μL of purified wCp149 capsids were applied to a glow-discharged UltrAuFoil^®^ R2/2 holey gold film grid and vitrified using a FEI Vitrobot™ (Mark III). The grids were imaged using FEI Titan Krios operated at 300 kV. Low-dose (~ 30 e^-^/Å^2^) cryo-EM images, at a nominal magnification of 81,000x (equivalent to 0.86 Å per pixel), were acquired on a Gatan K2 Summit direct electron detector using super resolution mode. Leginon automated data collection software [33] was used to collect 251 images. A total of 25,800 particles were semi-automatically extracted using e2boxer_old.py from EMAN2 [34]. Contrast transfer function parameters were estimated using ctffind4 [35]. All particles were subjected to reference-free 2D classification using Relion (v2.1) software [36]. Particles in classes that showed blurred density were discarded at this stage leaving 23,700 particles. A low-resolution initial model was built using e2initialmodel.py from EMAN2. This model was used as a starting model in Relion 3D auto-refine. The dataset was split into two parts for refinement, which was performed iteratively until the correlation between the two halves of the dataset converged. The resolution cutoff was determined using the gold standard Fourier shell correlation of 0.143 (Fig 7). A final 4.5Å 3D electron density map was obtained using 7,911 particles after applying a negative B-factor of - 214, obtained using the Guinier fitting procedure from Relion [37], to sharpen the density. The final density map of WHV capsid was deposited in the EMDB database as 9031.

### Capsid model determination

To model the WHV capsid structure, the crystal structure of wY132A with Y132A changed back to Y132 was rigid-body docked into a density map corresponding to a dimer that was extracted from the WHV capsid map. Using molecular dynamic flexible fitting (MDFF) implemented in VMD and NAMD [19, 38, 39], the mutated wY132A structure was fit into the capsid density. A WHV capsid structure was generated using the fitted wY132A structure and this was rigid-body fitted into the WHV capsid density. By applying symmetry restraints [40], the generated WHV capsid structure was fit into to WHV capsid density using MDFF. Molecular coordinates from the final frame of MDFF was extracted and refined using the PHENIX real space refinement function with imposed icosahedral symmetry [20] (S2 Table). The determined WHV capsid coordinates were deposited in the protein data bank as 6EDJ. All structures and maps were analyzed and displayed using UCSF Chimera [41].

## Acknowledgement

This research used resources of the Advanced Light Source, which is a DOE Office of Science User Facility under contract no. DE-AC02-05CH11231. We thank Jay Nix for his assistance during X-ray data collection at beamline 4.2.2. We also would like to acknowledge the Purdue cryo-EM facility, Thomas Klose, and Valerie Bowman for collecting cryo-EM images. We also acknowledge Alex Kukreja for his preliminary experiments with WHV and antiviral molecules. This work was supported by NIH R01-AI118933 to AZ and the Indiana Clinical and Translational Sciences Institute.

## Supporting information

**S1 Fig.**
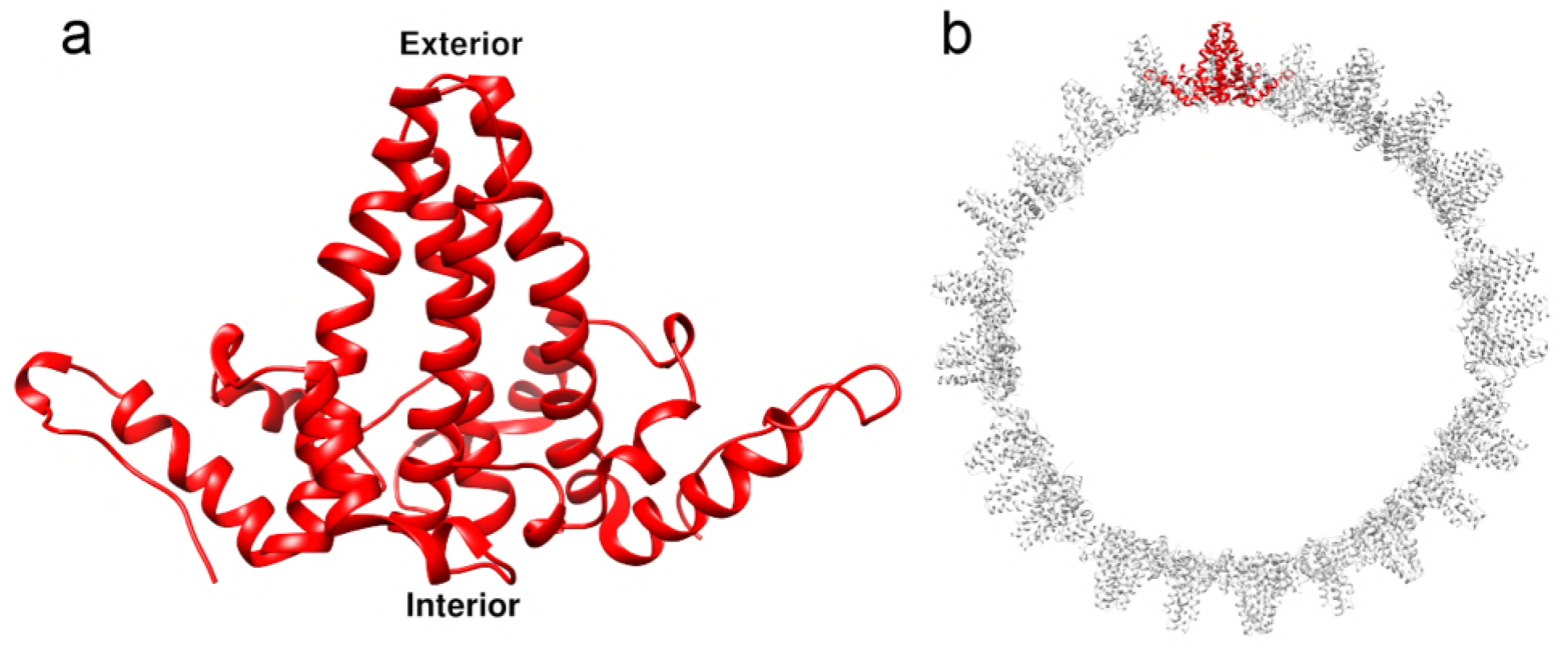
The arrangement of dimers in an HBV capsid. (a) One HBV dimer has the base facing the capsid interior and the spike tips protruding toward the capsid exterior. (b) One section of the HBV capsid showed the location and the arrangement of one HBV dimer (red) in the capsid.

**S2 Fig.**
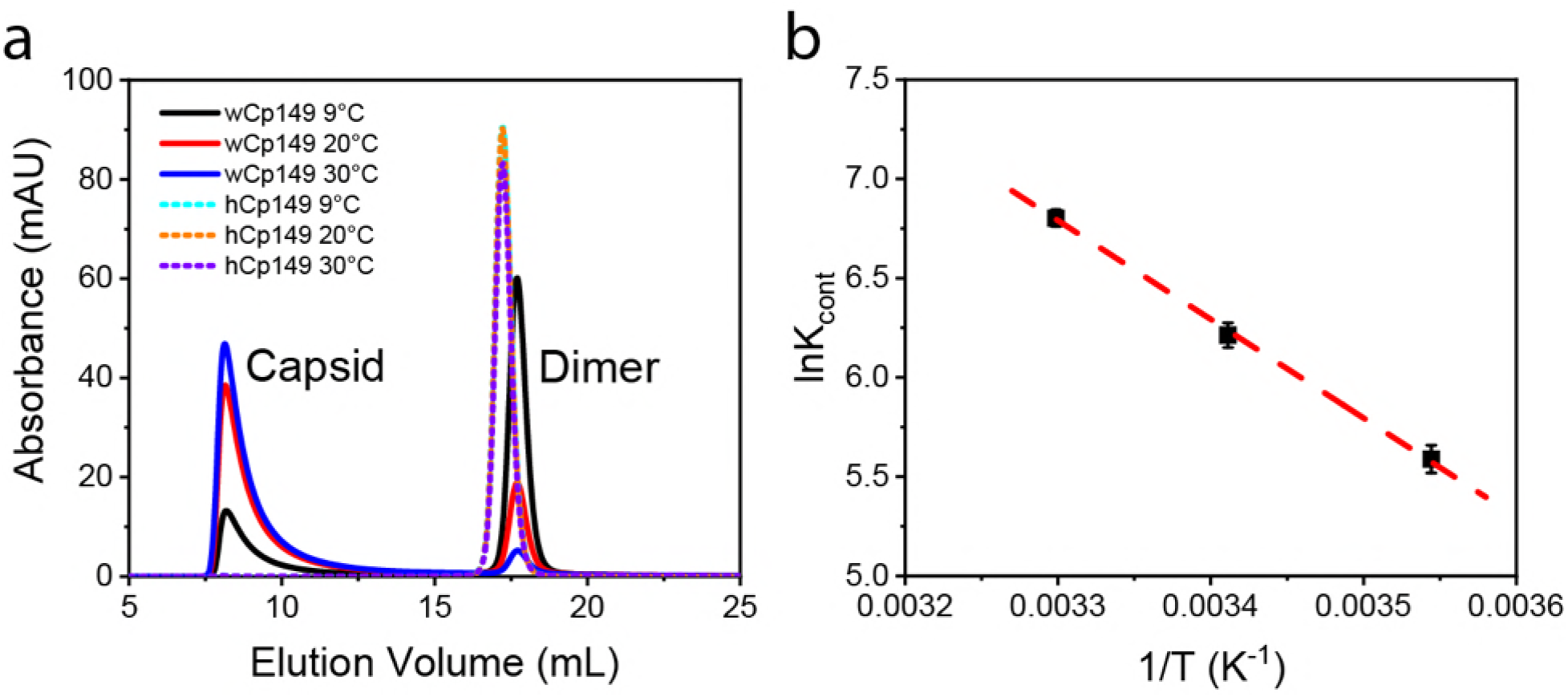
wCp149 assembly is thermodynamically favored. (a) wCp149 shows substantial assembly under various temperature, while hCp149 showed no assembly under the same assembly conditions. (b) van’t Hoff plot of wCp149 assembly shows that wCp149 assembly reaction has a linear temperature dependence in the assembly temperature range.

**S3 Fig.**
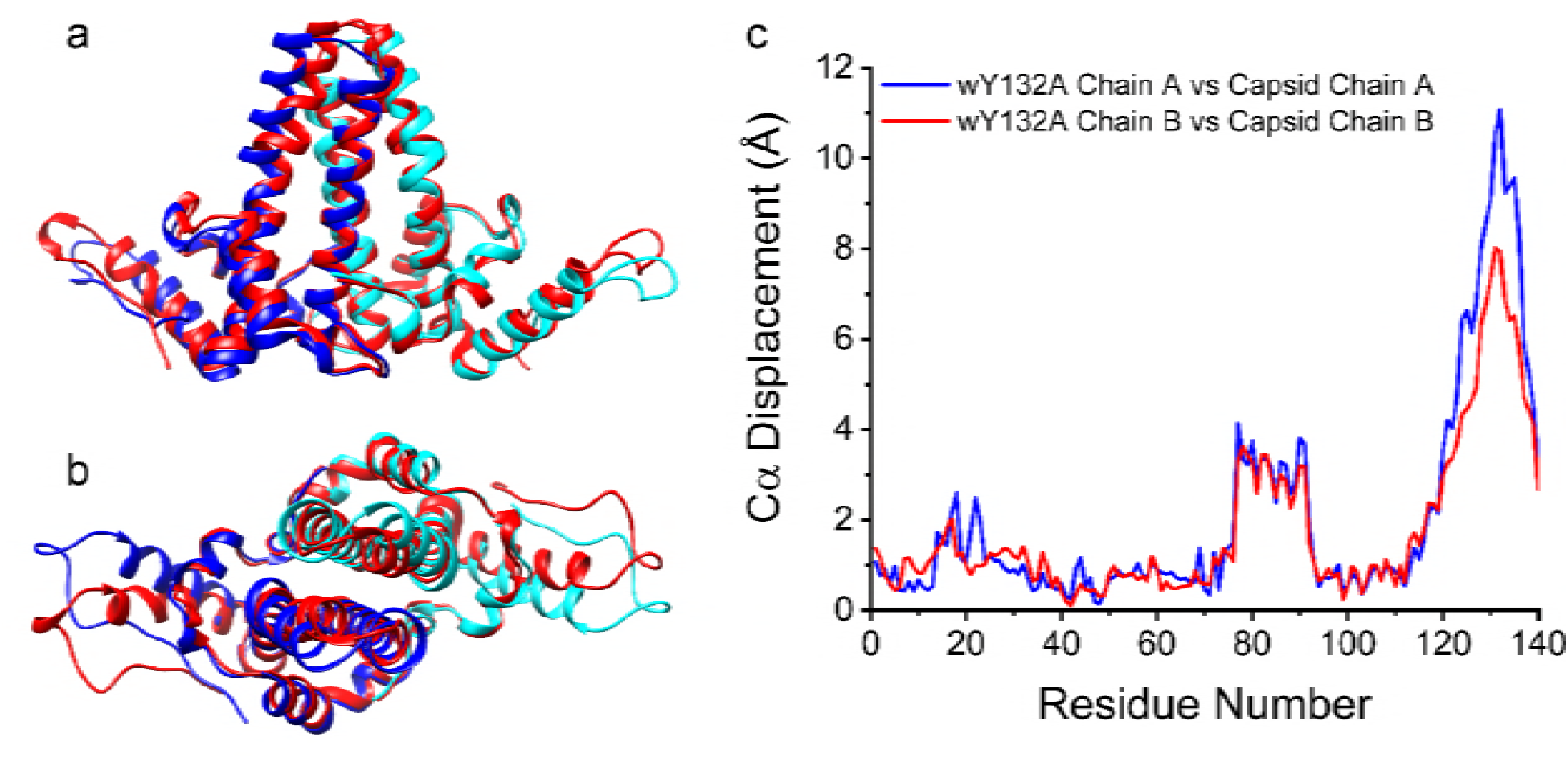
wY132A shows distinct structural differences compared to dimers from the HBV capsid. Side view of wY132A AB dimer overlaid with HBV AB dimer (red; PDB:1qgt) (a) shows that overall similar structures. Top view of wY132A overlaid with HBV AB (b) displays obvious structural changes at the α-helix 5 and C-termini. By calculating and plotting Cα displacement against residue numbers, wY132A AB show dramatic displacement comparing to HBV AB (c) near the spike tips (around 4 Å) and at the C-terminus (over 8 Å).

**S4 Fig.**
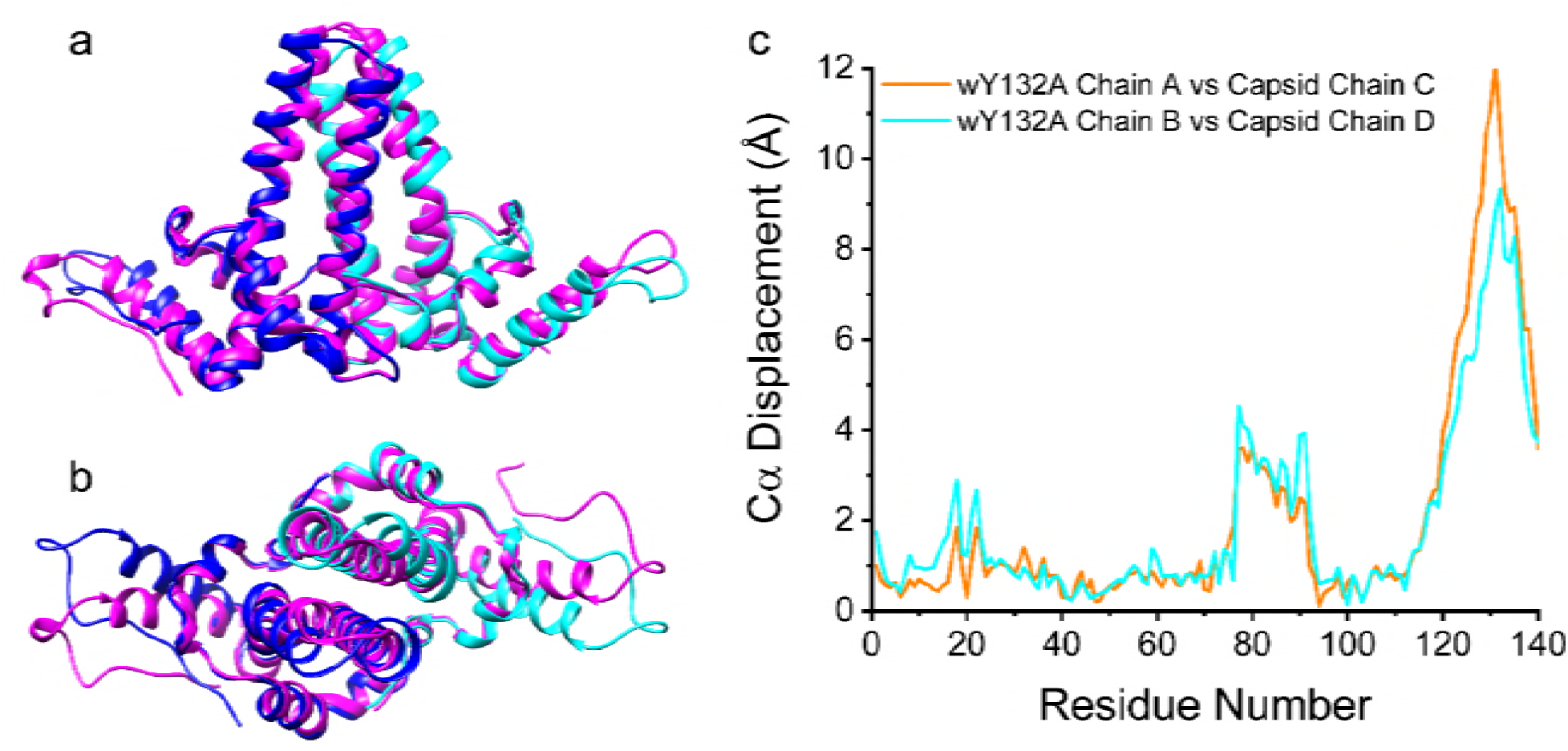
Side views of wY132A AB dimer overlaid with HBV CD dimer (hot pink; PDB:1qgt) (a) show that overall similar structures. Top views of wY132A overlaid with HBV CD (b) display obvious structural changes at the α-helix 5 and C-terminus. By calculating and plotting Cα displacement against residue numbers, wY132A AB show dramatic displacement compared to HBV CD (c) near the spike tips (around 4 Å) and at the C-terminus (over 10 Å).

**S1 Table.** Crystallographic data collection and refinement statistics.

|  | **wCp149-Y132A\*** |
| --- | --- |
| **PDB accession code** | 6ECS |
| **Wavelength (Å)** | 1 |
| **Resolution range (Å)** | 63.09 - 2.9 (3.004 - 2.9) |
| **Space group** | P 3<sub>2</sub> 2 1 |
| **Unit cell parameters** |  |
| a, b, c (Å) | 116.41 116.41 161.76 |
| α, β, γ (°) | 90 90 120 |
| **Total reflections** | 312472 (27830) |
| **Unique reflections** | 28665 (2817) |
| **Multiplicity** | 10.9 (9.9) |
| **Completeness (%)** | 99.93 (99.93) |
| **Mean I/sigma(I)** | 19.47 (1.28) |
| **Wilson B-factor** | 78.95 |
| **R-merge** | 0.1083 (1.638) |
| **R-meas** | 0.1136 (1.728) |
| **R-pim** | 0.0342 (0.5449) |
| **CC1/2** | 0.999 (0.589) |
| **CC\*** | 1 (0.861) |
| **Reflections used in refinement** | 28656 (2818) |
| **Reflections used for R-free** | 1993 (197) |
| **R-work** | 0.1984 (0.3492) |
| **R-free** | 0.2278 (0.3843) |
| **CC(work)** | 0.968 (0.696) |
| **CC(free)** | 0.964 (0.620) |
| **Number of non-hydrogen atoms** | 4393 |
| **Macromolecules** | 4393 |
| **Protein residues** | 542 |
| **RMSD bond length (Å)** | 0.007 |
| **RMSD bond angles (°)** | 0.86 |
| **Ramachandran favored (%)** | 96.23 |
| **Ramachandran allowed (%)** | 3.77 |
| **Ramachandran outliers (%)** | 0.00 |
| **Rotamer outliers (%)** | 0.00 |
| **Clashscore** | 9.71 |
| **Average B-factor (Å<sup>2</sup>)** | 84.23 |
| **Macromolecules (Å<sup>2</sup>)** | 84.23 |
| **Number of TLS groups** | 23 |

Statistics for the highest-resolution shell are shown in parentheses.

**S2 Table.**
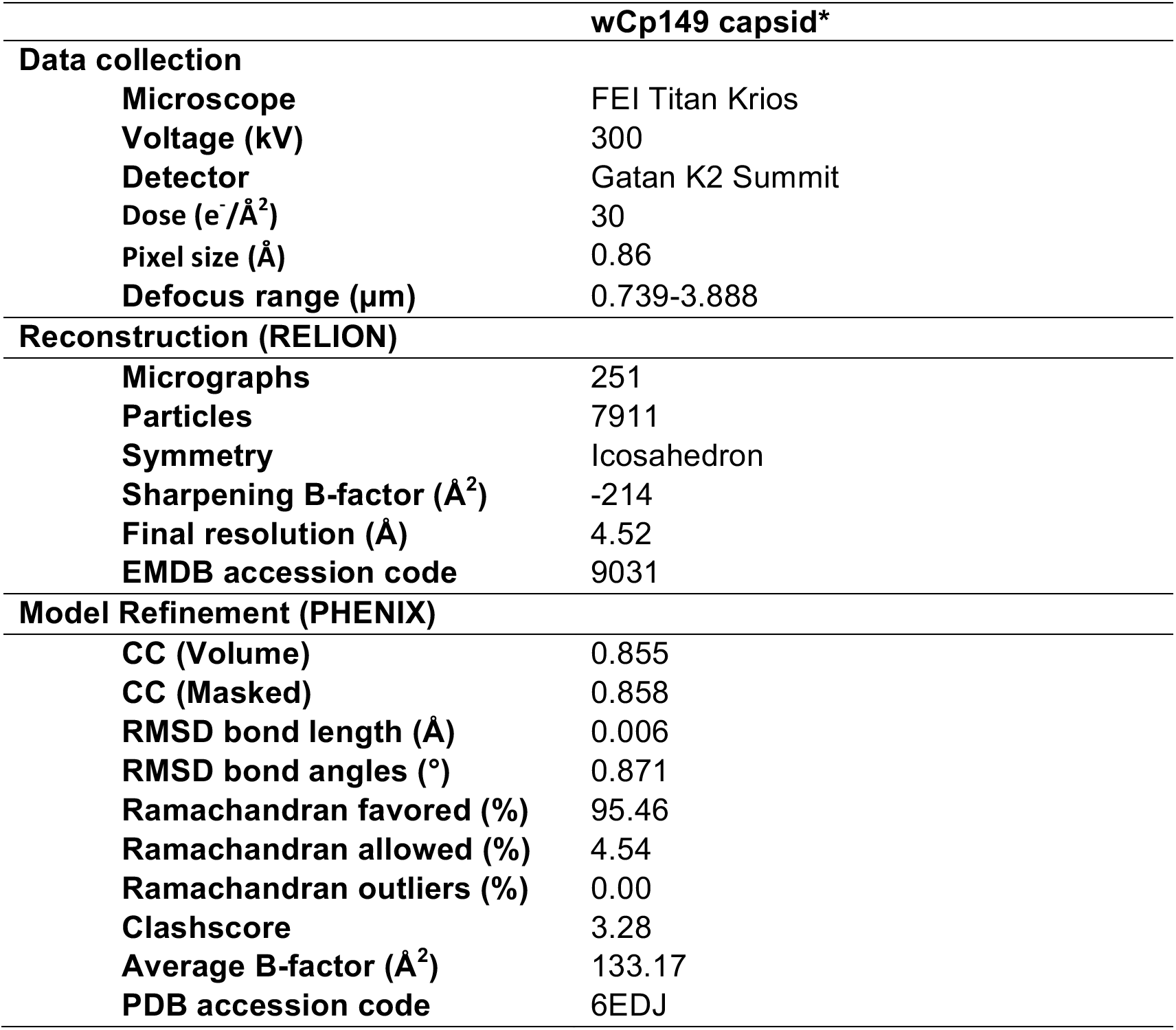
Cryo-EM data collection and refinement statistics.

